# Transgenerational developmental effects of immune priming in the red flour beetle *Tribolium castaneum*

**DOI:** 10.1101/422741

**Authors:** Nora KE Schulz, Marie Pauline Sell, Kevin Ferro, Nico Kleinhölting, Joachim Kurtz

**Affiliations:** Institute for Evolution and Biodiversity, University of Münster, Münster, Germany; Present address: Max Planck Institute for Chemical Ecology, Jena, Germany; Department of Entomology, University of Arizona, Tucson, USA

**Author notes:** Correspondence: Joachim Kurtz.

**Keywords:** innate immunity, immune priming, transgenerational effects, *Tribolium castaneum*, *Bacillus thuringiensis*, host parasite co-evolution, bacterial infection, oral infection

## Abstract

Immune priming, the increased chance to survive a secondary encounter with a pathogen, has been described for many invertebrate species, which lack the classical adaptive immune system of vertebrates. Priming can be specific even for closely related bacterial strains, last up to the entire lifespan of an individual, and in some species, it can also be transferred to the offspring and is then called transgenerational immune priming (TGIP). In the red flour beetle *Tribolium castaneum*, a pest of stored grains, TGIP has even been shown to be transferred paternally after injection of adult beetles with heat-killed *Bacillus thuringiensis*.

Here we studied whether TGIP in *T. castaneum* is also transferred to the second filial generation, whether it can also occur after oral and injection priming of larvae and whether it has effects on offspring development. We found that paternal priming with *B. thuringiensis* does not only protect the first but also the second offspring generation. Also, fitness costs of the immune priming became apparent, when the first filial generation produced fewer offspring. Furthermore, we used two different routes of exposure to prime larvae, either by injecting them with heat-killed bacteria or orally feeding them *B. thuringiensis* spore culture supernatant. Neither of the parental larval priming methods led to any direct benefits regarding offspring resistance. However, the injections slowed down development of the injected individuals, while oral priming with both a pathogenic and a non-pathogenic strain of *B. thuringiensis* delayed offspring development.

The long-lasting transgenerational nature of immune priming and its impact on offspring development indicate that potentially underlying epigenetic modifications might be stable over several generations. Therefore, this form of phenotypic plasticity might impact pest control and should be considered when using products of bacterial origin against insects.

## 1 Introduction

Over the last decade a wealth of new evidence has been put forward to demonstrate that invertebrate immune systems can possess forms of immune memory and are sometimes capable of highly specific responses (Cooper and Eleftherianos 2017; Milutinović and Kurtz 2016; Contreras-Garduño et al. 2016). The phenomenon enabling a stronger and faster immune response upon secondary infection has been termed immune priming and shows parallels in memory and specificity to trained immunity of vertebrates (Little and Kraaijeveld 2004; Kurtz and Armitage 2017; Kurtz 2005; Netea et al. 2011; Melillo et al. 2018). The trigger, specificity and duration of the priming can be extremely diverse. It has been shown that immune priming can be successful against bacteria (Roth et al. 2009), fungi (Fisher and Hajek 2015; Gálvez and Chapuisat 2014) and viruses (Tidbury, Pedersen, and Boots 2011). Immune priming can be achieved by introducing a sublethal dose of the parasite, an incapacitated, *e.g.* heat killed agent or using only specific molecules from the original pathogen, *e.g.* lipopolysaccharides (Milutinović and Kurtz 2016; Contreras-Garduño et al. 2016). Also, the route how the elicitor is introduced can vary, similar to differences in the route of infection in nature. For experiments involving priming, the priming agent is most commonly introduced via septic wounding and deposition into the haemocoel or orally via feeding (Milutinović and Kurtz 2016). Furthermore, also abiotic factors, e.g. thermal exposure have been shown to prompt this phenomenon (Wojda and Taszlow 2013; Eggert et al. 2015). Immune priming can be cross-reactive in some cases (Moret and Siva-Jothy 2003), while in others the host’s primed immune response can differentiate between bacterial species or even strains, mounting the best protection when the same pathogen is encountered twice (Roth et al. 2009; Medina Gomez et al. 2018).

Additionally, the duration of immune priming effects differs dramatically. In some cases, protection lasts across different life stages and throughout the entire life span of an individual (Thomas and Rudolf 2010; Khan et al. 2016; Pham et al. 2007). In some cases, the immune priming is even transferred to the offspring generation (Milutinović and Kurtz 2016; Roth et al. 2018; Dhinaut et al. 2018). This transgenerational immune priming (TGIP) can occur through either parent. While for the maternal side, the direct transfer of bacterial particles bound to egg-yolk protein vitellogenin has been shown to be involved in certain systems (Salmela et al. 2015), the detailed mechanistic underpinnings of immune priming in general and paternal TGIP in particular still remain to be discovered (Milutinović et al. 2016). It has been considered that epigenetic mechanisms, including DNA methylation, histone acetylation and miRNAs are involved (Vilcinskas 2016; Eggert et al. 2014).

As with any other immune response also the fitness costs of immune priming including those for storing the information have to be considered. These costs are not constraint to a direct reduction in fertility but can also become visible in delayed development or smaller body mass if the priming occurs before the organism reaches maturity. Furthermore, negative effects might only become visible in the offspring generation. In the Coleopteran, *Tenebrio molitor*, maternal priming prolonged offspring larval development (Zanchi et al. 2011) and the strength of this effect depended on the Gram type of the bacteria used for priming (Dhinaut et al. 2018). Immune priming beneficial to the mother can even increase offspring susceptibility to the same parasite (Vantaux et al. 2014). These are all factors demonstrating the complexity of immune priming and showing that this term probably covers several distinct (Pradeu and Du Pasquier 2018). It makes predicting host-parasite co-evolution and the emergence of resistance against bacterial pesticides much more difficult if we consider that several forms of immune priming can occur in the same species across different life stages and generations with different consequences.

Immune priming has intensively been studied in the red flour beetle *Tribolium castaneum*, which is a widely abundant pest of stored grains. In this beetle, immune priming has been demonstrated in different life stages, i.e. larvae and adults, as well as within and across generations (Milutinović et al. 2016). In this species, immune priming can be highly specific, down to the bacterial strain and can be passed on via both parents (Roth et al. 2009; Roth et al. 2010). Two different routes of priming and infection were used. Oral infections with spores only work in larvae and the protective benefits of priming with the supernatant of the spore culture have so far only been studied within generation, mostly even within life stage (Milutinović et al. 2014; Futo et al. 2017; Greenwood et al. 2017). Therefore, the effectiveness of the priming was only confirmed for a few days after exposure. The other priming and infection method uses vegetative cells, which are inactivated for the priming and are directly introduced into the body cavity via septic wounding (Tate et al. 2017; Khan et al. 2016; Milutinović et al. 2016). In this case, immune priming of adults can be transferred to their offspring and a protection against infection can still be observed in the adults of the offspring generation (Roth et al. 2010; Eggert et al. 2014). But, these different priming techniques and routes of infection lead to different responses as is evident in differential gene expression and immune system activity (Behrens et al. 2014). The pathogen used in most studies of priming in *T. castaneum* is the entomopathogenic and endospore forming bacterium *Bacillus thuringiensis* (Jurat-Fuentes and Jackson 2012). Proteins from *B. thuringiensis*, so-called Cry toxins are widely used for their insecticidal activity in transgenic crops (Lacey et al. 2015; Pardo-López et al. 2013). Therefore, the study of immune priming in this host parasite model system does not only advance basic research and our understanding of the invertebrate immune system but is also helpful for applied approaches and improving insect control strategies.

With our study we shed further light on the different forms of immune priming against *B. thuringiensis* that can be observed in *T. castaneum*. We here investigated the transgenerational effects caused by three different types of priming, *i.e*. priming by injection of larvae and male adults and oral priming of larvae by monitoring the development, fitness and survival of bacterial infection (challenge). As paternal TGIP so far has only been tested in the first offspring generation (Roth et al. 2010), we here expanded the experimental time frame to include the adult grandparental generation, investigating whether TGIP is a multigenerational phenomenon extended to more than one subsequent generation. Studies on larval priming have been mainly focused on within generation immune benefits (Milutinović et al. 2016). We therefore here wanted to investigate whether larval TGIP via the oral or septic wounding infection route exists and whether the offspring is affected in a different way by parental treatment.

## 2 Materials and Methods

### 2.1 Model organisms

Beetles were derived from a population originally collected in the wild in Croatia in 2010 (Milutinović et al. 2013). Until the start of the experiment, beetles were kept in populations of more than 2,000 individuals in plastic boxes with heat sterilized (75 °C) organic wheat flour (type 550) enriched with 5 % brewer’s yeast. Standard breeding conditions were set at 70 % humidity and 30 °C with a 12 h light/dark cycle.

In all priming treatments and infections, different entomopathogenic gram-positive *B. thuringiensis* strains were used. *B. thuringiensis* and its Cry toxins are widely used as insecticides and together with *T. castaneum* form a well-established system to study host parasite co-evolution (Roth et al. 2009; Milutinović et al. 2013; Contreras et al. 2013; Pardo-López et al. 2013). For priming and challenge by injection we used vegetative cells from *B. thuringiensis* (*Bt*) strain DSM 2046 (German Collection of Microorganisms and Cell Cultures, DSMZ). For the treatments concerning priming and infection by oral uptake, spores and supernatant from *Bt morrisoni bv. tenebrionis* spore cultures *(Btt*, Bacillus Genetic Stock Center, Ohio State University, Ohio, USA) were used. Additionally, *Bt407cry–* (*Bt407*, kindly provided by Dr. Christina Nielsen-Leroux, Institute National de la Recherche Agronomique, La Minière, 78285 Guyancourt Cedex, France) served as a negative control in the oral priming experiment, as it does not produce Cry toxins and does not lead to immune priming nor mortality upon ingestion (Milutinović et al. 2013; Milutinović et al. 2014).

### 2.2 TGIP in adults

In this experiment we wanted to investigate, whether paternal TGIP persists past the first filial (F_1_) generation (Roth et al. 2010; Eggert et al. 2014) and therefore provides survival advantages upon *Bt* infection to the second filial (F_2_) generation. Additionally, we measured the fertility of the primed males and their offspring to determine potential costs of TGIP. For an overview of the experimental design see Figure 1.

**Figure 1.**
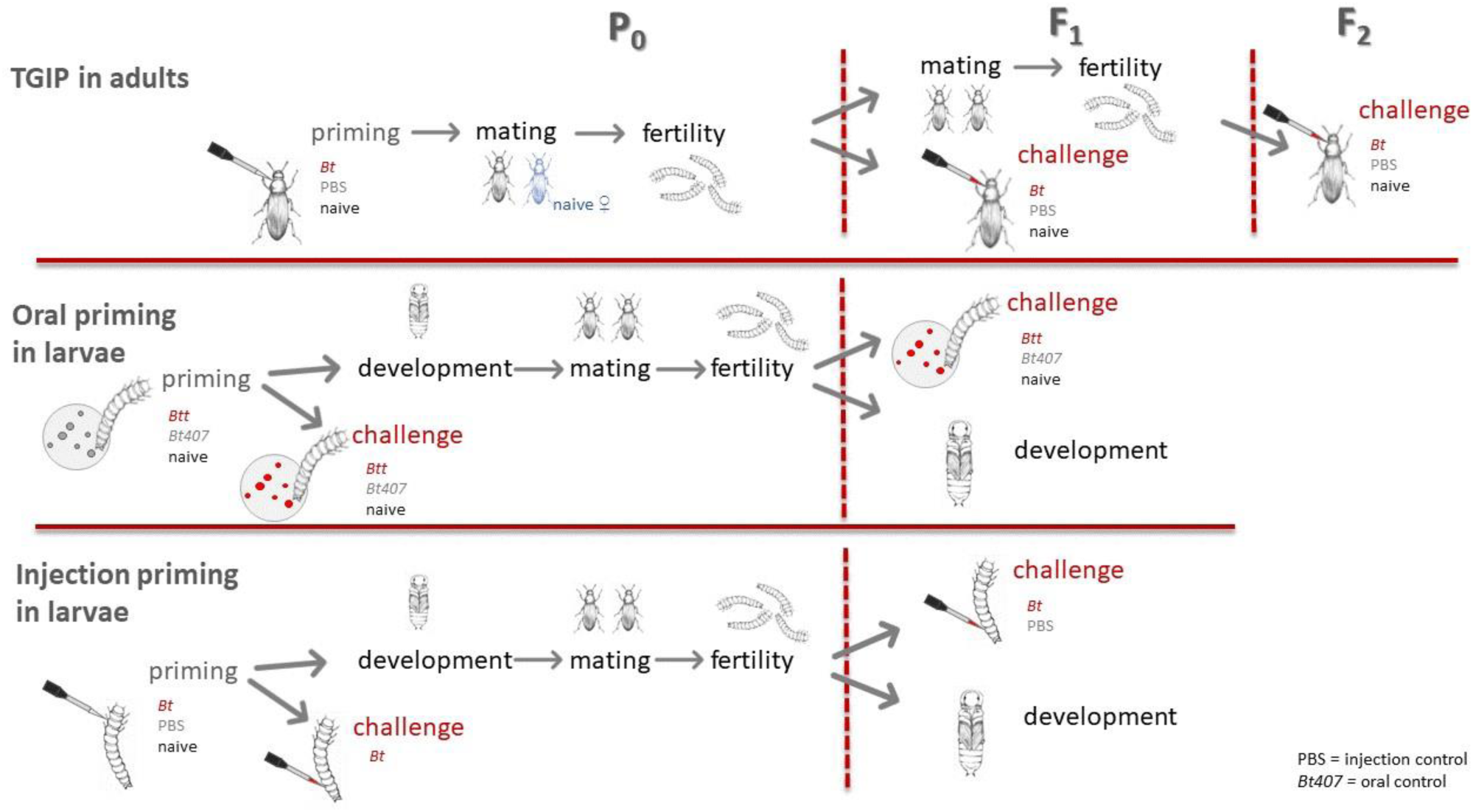
Overview of the experimental design

#### 2.2.1 Priming of the parental (P_0_) generation

To set up the P_0_ generation for this experiment around 2000 beetles from a general stock population were put into a plastic box containing 250 g of flour with yeast. After an oviposition period of 24 h the adults were sieved off and put into a new box for a second 24 h oviposition period. When the offspring had reached the pupal stage, their sex was determined, and all beetles were kept individually from here on onwards.

For the priming injections one week after eclosion, 60 male adults were either injected with heat-killed *Bt* suspended in PBS at a concentration of 1*10^9^ cells per ml, PBS only to control for the wounding or left naïve. The priming suspension was directly injected into the dorsal vessel by dorsally puncturing the epidermis between head and pronotum in a flat angle to minimize tissue damage. Heat-killed *Bt* were produced from an overnight culture as previously described (Roth et al. 2009; Ferro et al. 2017). A nanoinjector (Drummond Nanoject II) was used for this procedure with the injection volume set to 18.4 nl (∼20,000 cells per injection in the *Bt* treatment). Survival after the priming procedure was recorded 24 h later.

#### 2.2.2 Mating and fitness of P_0_ and F_1_ generation

Single mating pairs with naïve, virgin females were set up (n=39-57). Mating pairs were kept in plastic vials containing 6 g of flour and left to lay eggs for two consecutive three-day long oviposition periods. Thirteen days after the end of the respective oviposition period, larvae were counted for each pair and individualized into 96 well plates with flour. For the analysis, data from both oviposition periods were combined.

The sex of the offspring was determined when they had reached the pupal stage. One female and one male offspring from each single pair formed a new mating pair to produce the F_2_ generation, leading to mating of full siblings (n=29-53). Mating, oviposition and individualization of offspring larvae were carried out in the same way as described for the parental generation with the exception of the oviposition periods being shortened to 24 h. The fertility of F_1_ pairs was recorded as live larvae 12 days post oviposition (dpo).

#### 2.2.3 Bacterial Challenge of adults of F_1_ and F_2_ generation

The priming of adult males of *T. castaneum* with heat-killed *Bt* leads to an increased survival rate in their adult offspring when infected with a potentially lethal dose of the same bacteria (Eggert et al. 2014). Whether this phenomenon is also transferred to subsequent generations has so far not been investigated. We therefore exposed individuals of the F_1_ and F_2_ generation to a bacterial challenge after the P_0_ generation had received a priming treatment. Bacteria were cultured, washed and their concentration in PBS adjusted as for the priming procedure without the heat-killing step (2.2.1). One week after eclosion animals of both sexes were injected with a volume of 18.4 nl. The injection either contained *Bt* cells at a concentration 10^7^ vegetative bacterial cells per ml (∼200 cells per injection) in PBS or only PBS as a control and was performed in the same manner as described for priming (2.2.1). A second control consisted of a naïve group that received no injections. In the F_1_ generation, three adult siblings from each family were used, one for each challenge treatment (n=31-44). This was the same for the F_2_ generation, but here the challenge was performed on adults originating from two consecutive ovipositions of the same families (oviposition 1: n=16-42, oviposition 2: n=24-45). Injections were carried out in the same manner as for the priming treatment (2.2.1). Afterwards, the beetles were kept in individual glass vials and their survival was recorded 24 h post challenge.

### 2.3 Transgenerational effects of larval priming

Within generation immune priming of *T. castaneum* larvae with *B. thuringiensis* can be achieved by two different infection routes, *i.e.* oral ingestion of spore conditioned supernatant or by the introduction of heat-killed vegetative cells into the hemolymph by pricking with a needle dipped in bacterial solution or direct injection (Behrens et al. 2014; Ferro et al. 2017). Here, we investigated whether priming via either of these infection routes affects fertility. For this, larvae from a 24 h oviposition period of our stock population were exposed to one of two oral treatments, one of two injection treatments or left naïve (for priming treatment details see 2.3.1 and 2.3.2). During the pupal stage the sex of the individuals was determined and once they had reached sexual maturity, single mating pairs were formed within each priming treatment (n=57-66). Pairs were allowed to mate and produce eggs for two consecutive 24 h oviposition periods. Afterwards, the adults were sieved off and offspring larvae were counted 14 dpo. For the analysis, data from both oviposition periods were combined.

Furthermore, we wanted to know whether the oral or injection immune priming of larvae can also be transferred to the F_1_ generation, as has been observed in the priming of adult *T. castaneum* (Tate et al. 2017; Roth et al. 2010). To answer this, the following TGIP experiments on larvae were conducted applying oral and injection immune priming and challenge protocols. For an overview of the experimental design see Figure 1.

#### 2.3.1 Oral priming and challenge of larvae

For the culturing and sporulation of *B. thuringiensis* we followed the method given in Milutinović *et al.* (2013). Milutinović *et al.* (2014) describe the methodology to orally prime larvae with *Bt* spore supernatant. In short, for the oral priming and challenge the spore supernatant or spores are provided to the beetle by mixing them with flour and PBS, pipetting the mixture into a 96 well plate and letting the diet dry to form flour discs. In addition to the *Btt* treatment, *Bt407* was used as a negative control in the priming and challenge procedure because the supernatant from its spore culture does not provide a priming effect nor do its spores cause mortality upon ingestion (Milutinović et al. 2014; Milutinović et al. 2013). As a third group a naive control was included with pure PBS to produce the flour discs.

The P_0_ generation originated from approximately 1000 beetles from our stock population ovipositing for 24 h. Larvae of the P_0_ generation were exposed to the priming diets 14 dpo for 24 h (n=320). After the priming, a subgroup of the primed larvae was transferred onto naïve flour discs, on which they remained until the oral challenge. The within generation challenge was performed to confirm successful priming. The challenge took place 19 dpo, *i.e.* five days after the exposure to the priming diet, in a full factorial design. Besides the challenge diet of *Btt* spores, two controls were included using either *Bt407* spores or flour discs prepared with pure PBS (n=40). The spore concentration was adjusted to 5*10^9^ spores per mL. Larvae stayed on their respective flour discs for the rest of the experiment. Survival after challenge was recorded daily for the next eight days.

As mentioned above, a subgroup of the F_1_ generation was orally challenged as well. This group was produced by the mating of single pairs coming from the same priming group. One individual from each mating pair was used for each of the three challenge treatments (n=71-76). The challenge was conducted in a similar manner as for the P_0_ generation, but without the naïve control. Instead it included two different spore concentrations to counteract the possibility of too high or too low mortality rates. The spore concentration was set to either 1*10^10^ spores per ml (high dose) or 5*10^9^ spores per ml (low dose). Larvae were put on naïve flour discs at 14 dpo to ensure similar development as in the P_0_ generation and avoid early pupation, as the development in lose flour is considerably faster than on flour discs. The challenge took place 19 dpo and again survival was monitored for eight days.

To determine potential costs of the immune priming, we monitored the development of the remaining individuals of the P_0_ generation after priming that were not used in the challenge (*Btt* and *Bt407*: n=196, naïve: n=280) and their offspring produced from the mating pairs, which came from the same priming treatment (70-75). In the P_0_, pupation rates were monitored between 21 dpo and 25 dpo and the proportion of eclosed adults was recorded 27 dpo. The offspring larvae were individualized 14 dpo and kept in lose flour the entire time. They were checked for pupation between 19 dpo and 23 dpo and their eclosion rates were noted 28 dpo.

#### 2.3.2 Priming and challenge of larvae by bacterial injection

Priming by injection of heat killed *Bt* cells took place 14 dpo. The larvae for this experiment came from a 24 h oviposition of ∼1000 beetles from our stock population. The procedure also included an injection control in which only PBS was used and a naïve group (n=244). Heat-killed priming bacteria were produced as described above (2.2.1). Priming injections had a volume of 18.4 nl and were placed in a flat-angle laterally under the epidermis of the third-last segment using a nanoinjector (Drummond Nanoject II). The bacterial concentration was adjusted to 1*10^9^ cells per ml (∼20,000 cells per larvae). After the injection, larvae were kept individually in 96 well plates containing flour.

We performed a within generation injection challenge to confirm the success of the priming. During the bacterial challenge 19 dpo, *i.e.* five days post priming a subgroup of the animals was injected with 18.4 nl of either vegetative *Bt* cells at a concentration of 1*10^7^ cells per ml suspended in PBS or only PBS (n=48). Challenge injections were placed in the dorsal vessel at a flat angle dorsally under the epidermis of the first thoracic segment to minimize tissue damage. After the challenge injection, larvae were continued to be kept individually, and their survival was checked seven days later. This challenge procedure was performed on a subgroup of the F_1_ generation in the same manner, which was produced from single pairs within the same priming group, which produced eggs for two consecutive 24 h periods (n=96). Again, survival was measured after seven days.

Also, for the injection priming, we wanted to test whether the treatment was costly and impacted the development. We therefore checked the proportion of pupae in a subgroup of the P_0_ generation (n=196) 23 dpo and the proportion of eclosed adults in the F_1_ generation (n=72-103) 27 dpo. The F_1_ generation was produced from single mating pairs within a priming treatment and offspring larvae were individualized 14 dpo, *i.e.* the age their parents had been primed.

### 2.4 Statistical analysis

All statistical analyses were performed in R (R Development Core Team 2008) using RStudio (RStudio Team 2015). Additional packages utilized included: MASS (Venables and Ripley 2002), lme4 (Bates et al. 2015), multcomp (Hothorn et al. 2008) and survival (Therneau and Grambsch 2000). Data concerning larval survival and development until pupation were tested in a Cox proportional hazard analysis, after it had been ensured that the assumption of hazards being proportional over time had been fulfilled. When this was not the case, generalized linear mixed effects models (GLMM) with a binomial distribution and experimental block as random factor were applied on data for one specific time point for pupation rates. This method was also used to examine eclosion rate. Tukey Honest Difference (THD) was applied *post hoc* to determine significant differences between individual treatment groups, while adjusting the p-values for multiple testing. *X*^2^-tests were used to analyze survival after injection challenge in cases for which random factors did not apply.

## 3 Results

### 3.1 Adult TGIP is transmitted to the F_2_ generation

We first wanted to confirm successful TGIP in the adults of the F_1_ generation. Due to an unusually high death rate in the beetles injected only with buffer, we did not observe significant differences in mortality between the beetles exposed to bacteria (challenge) and the injection control regardless of paternal priming (N=232, *X*^2^=0.707, p=0.4; Figure 2a). Within the *Bt* challenged group, there was a tendency towards effective TGIP, as we observed a trend towards increased survival in the group that had received the paternal priming treatment compared to the priming injection control (N=69, *X*^2^=3.401, p=0.065; Figure 2a). We did not see any difference between the priming treatments in response to challenge injection control (N=119, *X*^2^=0.473, p=0.78; Figure 2a).

**Figure 2:**
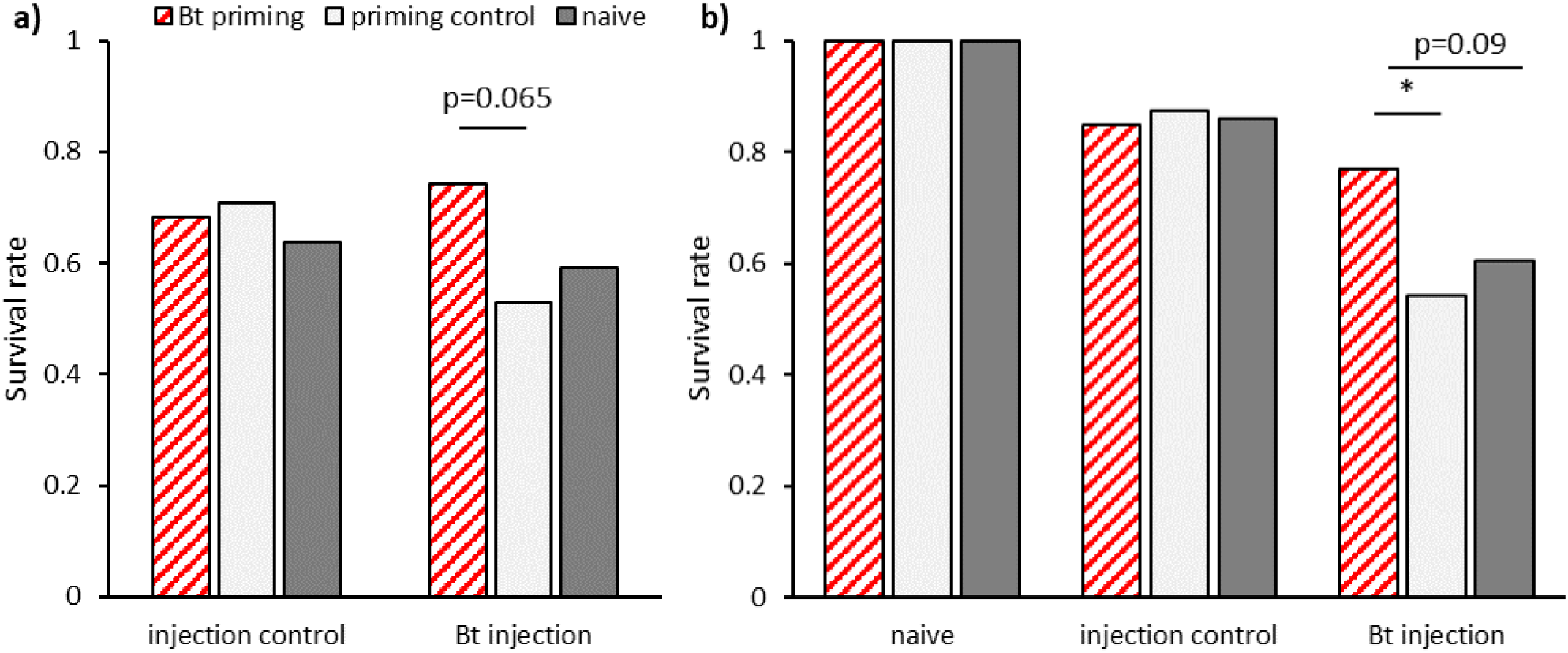
Survival of bacterial challenge after paternal TGIP. a) Adult F_1_ generation after parental priming (n=31-44) and b) adult F_2_ generation after grandparental priming (two experimental blocks: n=16-42 and n=24-45). Asterisks indicate significant differences at p < 0.05.

We then tested whether TGIP is also passed on to the successive generation. The challenge of the adult F_2_ generation proved to be effective, as significantly more beetles died after injection with live bacteria than of those that received control injections (GLMM: df=1, *X*^2^=23, p<0.001; Figure 2b). Furthermore, offspring, whose grandfathers had received a priming injection with heat-killed bacteria survived significantly better than those from the priming control group (GLMM: df=2, *X*^2^=7.3, p<0.05; THD: z=-2.492, p<0.05; Figure 2b). Survival in the naive control did not differ significantly from the priming control (THD: z=-0.827, p=0.68), but there was a tendency towards higher mortality compared to the offspring of primed grandfathers (THD: z=-2.090, p=0.09; Figure 2b). Therefore, the previously described TGIP in *T. castaneum* is transmitted past the first offspring generation at a comparable strength to the F_2_ generation.

We investigated possible costs of TGIP by counting live offspring two weeks after mating as a measure of reproductive success in the P_0_ and F_1_ generations. We could not observe any effect of parental priming treatment on fertility for the P_0_ (GLM: df=2, *X*^2^=3.399, p=0.18; Figure 3a) nor the F_1_ generation (GLM: df=2, *X*^2^=7.19, p<0.05; THD: priming control z=-0.527, p=0.86; naïve z=2.014, p=0.11, Figure 3b). However, the paternal priming control treatment significantly reduced fertility in the F_1_ generation and led to significantly less F_2_ larvae compared to the naïve control (THD: z= −2.381, p<0.05; Figure 3b). Therefore, paternal septic wounding but not paternal bacterial priming reduces the fitness of the F_1_ generation.

**Figure 3:**
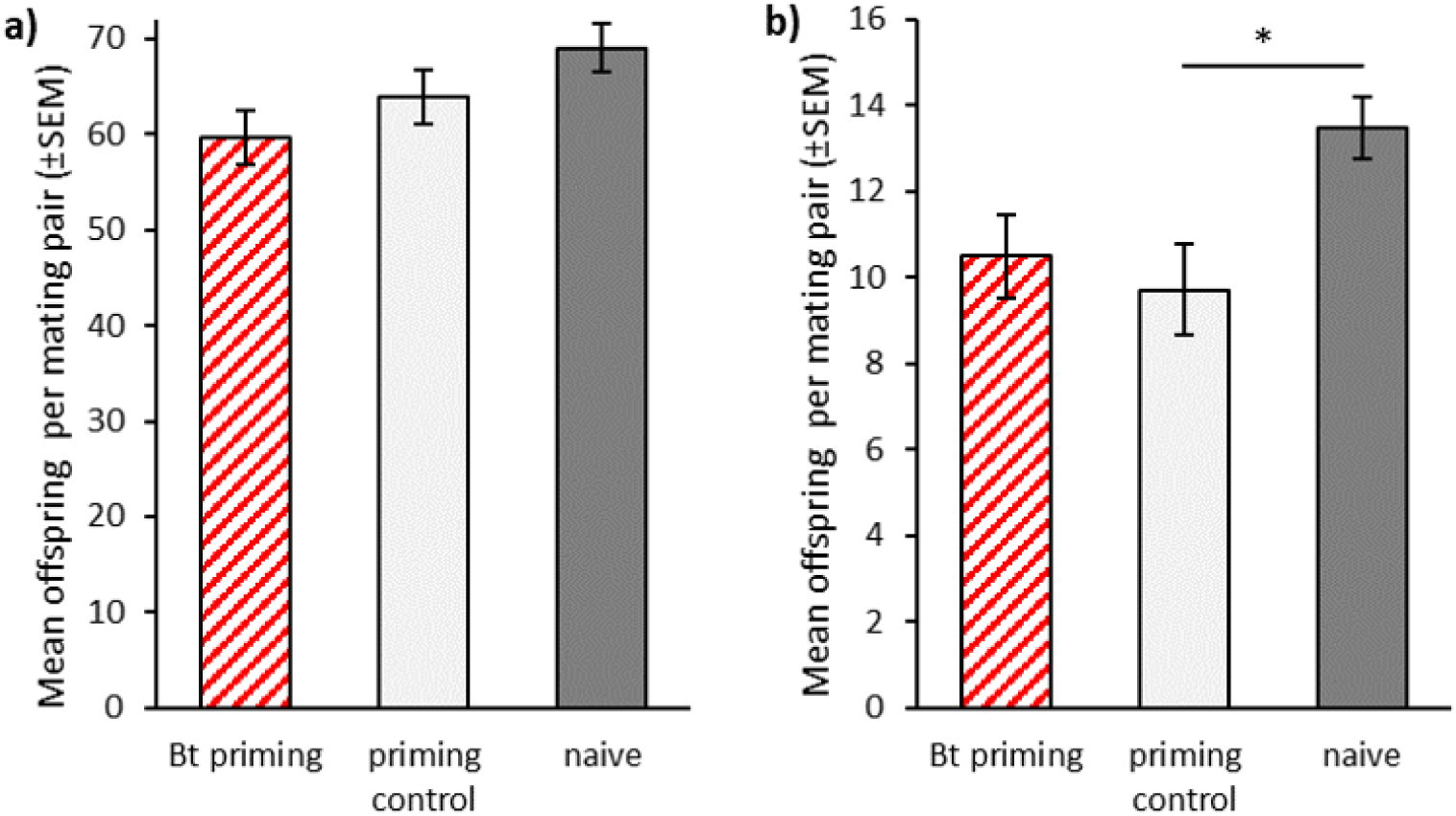
Male fertility after injection priming of adults. a) Mean offspring produced by primed males within 6 days (n=39-57) and b) mean offspring produced by offspring of primed males within 48 h (n=29-53). Asterisk indicates significant differences at p < 0.05

### 3.2 Transgenerational effects of priming in larvae

#### 3.2.1 Larval priming does not affect fertility

Neither oral nor injection priming of larvae with spore supernatant or heat-killed bacteria, respectively, significantly affected fertility compared to the control groups or the naïve individuals (GLM: df=4, *X*^2^=2.11, p=0.71, Figure S1).

#### 3.2.2 Oral priming affects development differently in treated and offspring generation

We monitored larval development after oral priming to discover potential additional costs and benefits of this treatment besides changes in survival rate upon infection. In the treated parental generation, there were significant differences in the pupation rates 21 dpo to 25 dpo (**Figure 4**a). Larvae treated with *Bt407* supernatant, a bacterial strain that has been shown to not cause any immune priming (Milutinović et al. 2014) and therefore served as a priming control, reached pupation faster than the *Btt* primed group (z=-2.906, p=0.0102). There also was a trend towards earlier pupation of the *Bt407* treated larvae compared to the naïve control (z=-2.28, p=0.059), while the *Btt* primed group and naïve control did not differ (z=-0.875, p=0.65). Additionally, there were differences in time until adult eclosion (GLMM: df=2, *X*^2^= 17.52, p<0.001;

**Figure 4.**
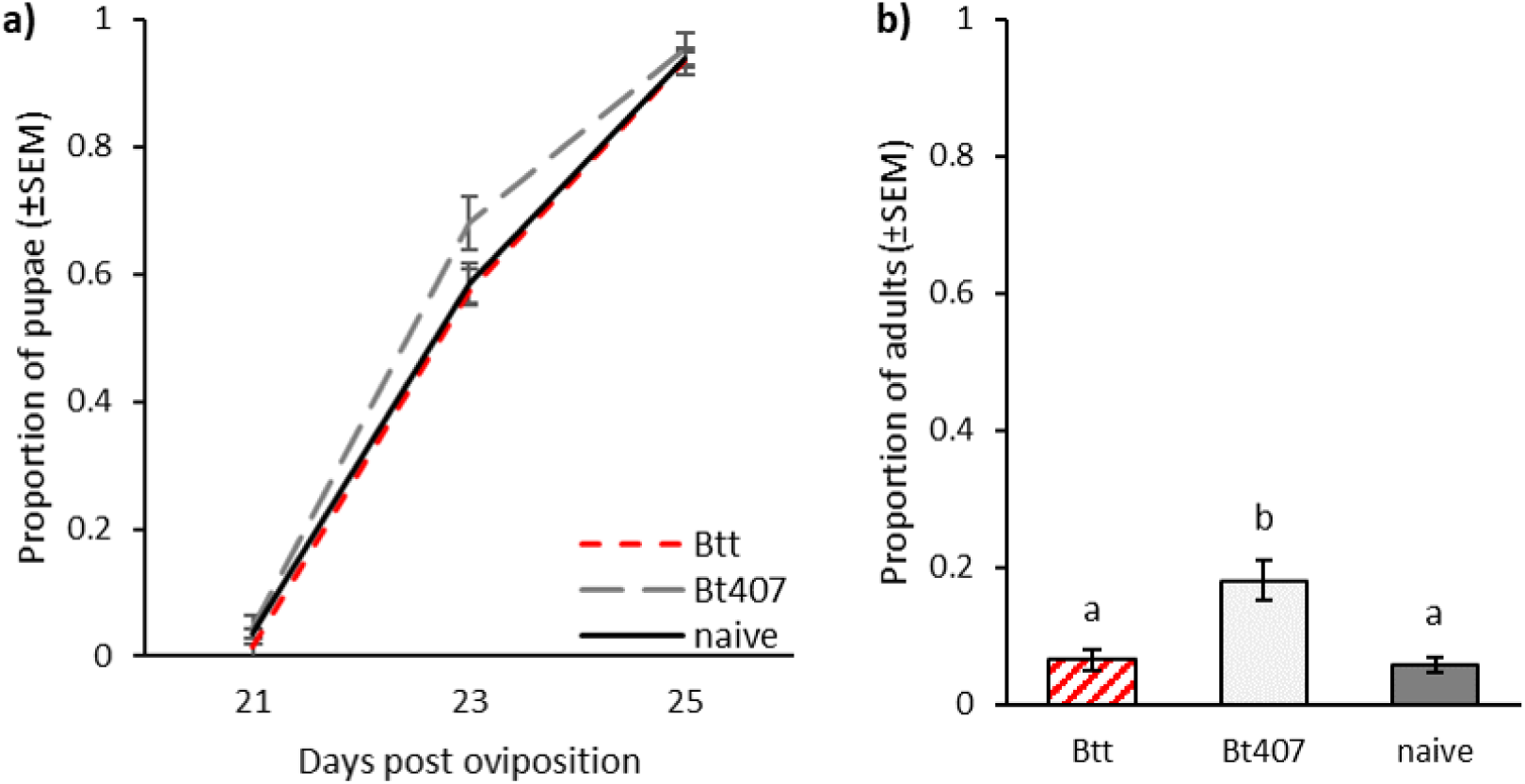
Development after oral priming during larval stage (n=196-280). a) Pupation rate for nine replicates b) Proportion of eclosed adults 28 dpo for nine replicates. Different letters indicate significant differences at p<0.05

**Figure 4**b). At 28 dpo significantly more pupae from the *Bt407* priming control had eclosed than from the *Btt* primed group (z= 2.98, p=0.008) and the naïve control (z= 3.802, p<0.001). Again, there was no difference between the *Btt* primed and naïve control (z= 0.569, p=0.84).

We also observed the development in the F_1_ generation to see if this was influenced by the parental oral priming. Larvae, whose parents were exposed to spore culture supernatant from *Btt* or *Bt407* developed significantly slower than offspring of the naïve control (GLMM: df=2, *X*^2^= 16.14, p<0.001; *Bt407*: z=3.83, p=0.002; *Btt*: z=3.832, p<0.001, Figure 5a). We found a similar effect for the development until adult eclosion, which on average was reached earliest by the naïve group (GLMM: df=2, *X*^2^= 14.17, p<0.001; *Bt407*: z=-3.213, p=0.004; *Btt*: z=-3.199, p=0.004; Figure 5b)

**Figure 5:**
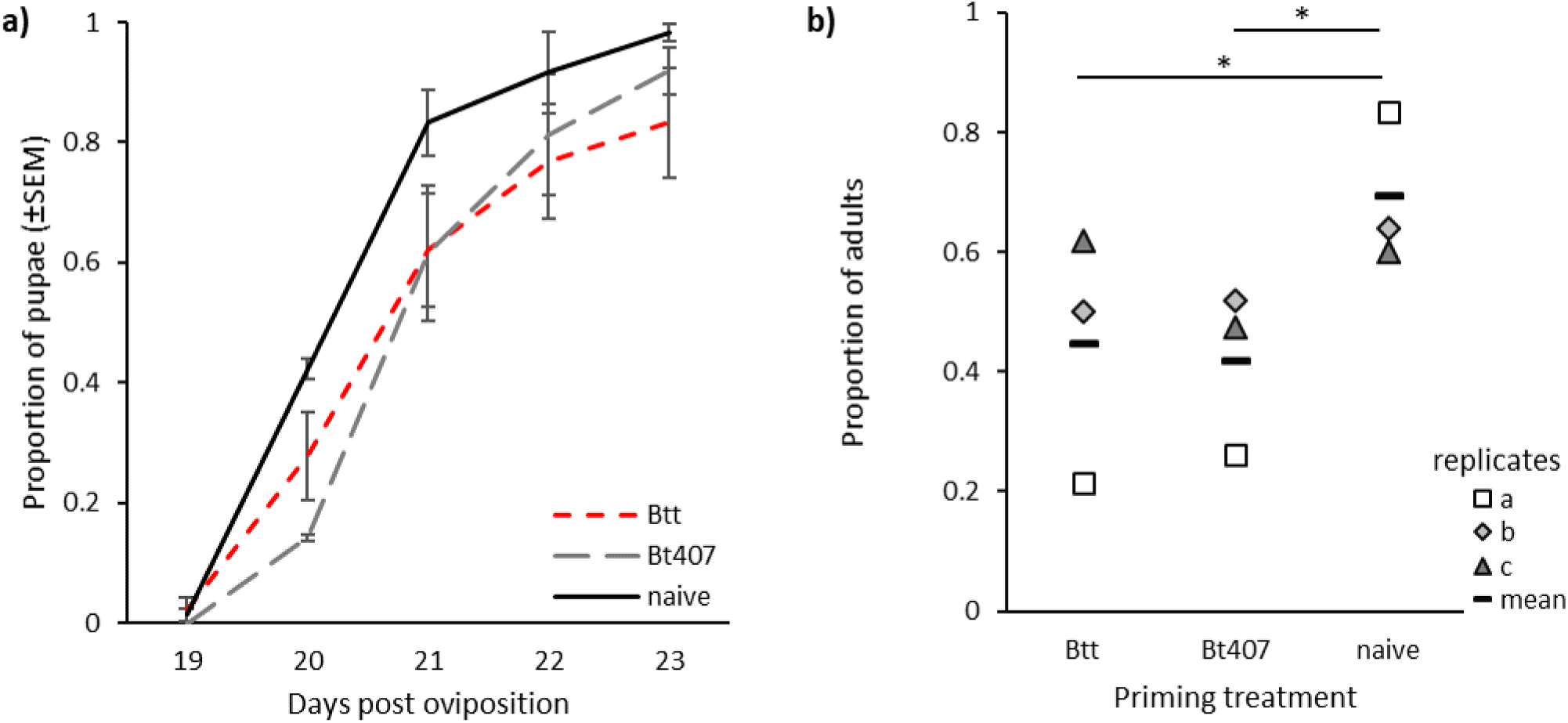
Offspring development after parental oral priming during the larval stage (n=70-75). a) Pupation rate b) Proportion of eclosed adults 28 dpo for three replicates. Asterisks indicate significant differences at p<0.05

#### 3.2.3 No survival benefits of oral TGIP for offspring generation

To test whether the exposure to spore supernatants led to a trans-generational priming effect, *i.e.* increased offspring survival upon infection, larvae of the primed P_0_ and the F_1_ generation were orally exposed to spores. In the primed P_0_ generation, the challenge with *Btt* spores killed the larvae at a significantly higher rate than the exposure to spores of *Bt407*, which served as the treatment control (df=1, *X*^2^=12.76, p<0.001; Figure S2). This, however was regardless of priming treatment, which did not lead to any significant differences (df=2, *X*^2^=0.63, p=0.73; Figure S2). This might be attributed to overall relatively low mortality rate after challenge with only 10.8 % of all exposed larvae dying. This probably was caused by the rearing of larvae in lose flour instead of flour discs for the period between priming and challenge, because of which many larvae might have already had reached a pre-pupal stage and stopped feeding.

Although mortality was higher, results for the offspring generation were similar (Figure S3). Again, the bacterial challenge proved to cause significant mortality at high (df=1, *X*^2^=96.63, p<0.001) and low concentration of spores (df=1, *X*^2^=47.1, p<0.001). Furthermore, survival depended on *Btt* spore concentration as the higher dose led to significantly higher mortality (df=1, *X*^2^=10.85, p<0.001). But, no effect of parental priming was observed (df=2, *X*^2^=0.69, p=0.71; Figure S3).

#### 3.2.4 Transgenerational effects of injection priming in larvae

In this part of the experiment we investigated potential effects of priming of larvae by injection on their development and the development of their offspring. Nine days after the priming, significantly less individuals from the control injection treatment had pupated compared to the naïve control (*X*^2^=8.466, p=0.003, Figure 6a). The addition of heat-killed bacteria to the injection reduced this effect, resulting in only a trend towards later pupation in the *Bt* priming treatment compared to the naïve control (*X*^2^=3.74, p=0.053, Figure 6a). There was no significant difference in the pupation rate between the *Bt* primed individuals and the injection control (*X*^2^=1, p=0.317, Figure 6a). In the F_1_ offspring generation we did not observe any effect of parental priming on the developmental speed, as the eclosion rate was similar for all treatment groups at 27 dpo (GLMM: df=2, *X*^2^=4.62, p=0.1).

**Figure 6:**
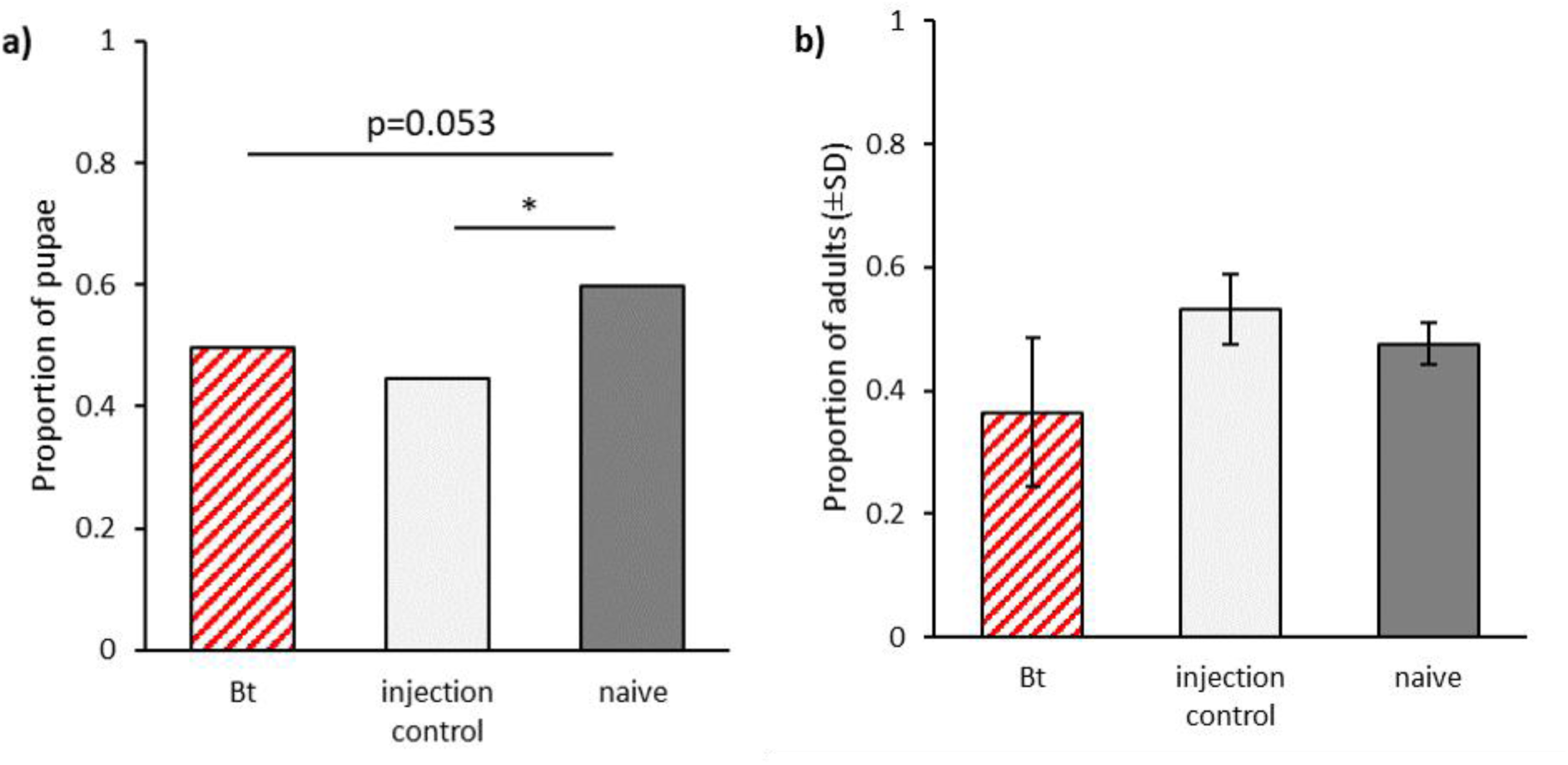
Development after parental injection priming during the larval stage. a) Proportion of pupae for P_0_ 23 dpo (n=196). b) Proportion of eclosed adults for F_1_ 27 dpo for two experimental blocks (n=72-103). Asterisks indicate significant differences at p < 0.05.

We challenged the parental and offspring generation by injecting a potentially lethal dose of *Bt* at 19 dpo, i.e. five days after the priming procedures for the parental generation. As the majority of mortality occurred within 24 hours of the bacterial injection, we did not use survival curves in the analysis, but instead used the survival rate differences after seven days for our analysis.

In the P_0_ generation priming did not lead to differential survival after the injection challenge, which caused between 23 % and 27 % mortality (df=2, *X*^2^=0.291, p=0.86). Finally, in the F_1_ offspring generation, the bacterial injection challenge caused significantly higher mortality than the injection control (GLMM: df=1, *X*^2^=244, p<0.001, Figure S4). However, also in this case parental priming did not significantly impact survival as there were no significant differences in mortality rates between the parentally primed group and the two controls (GLMM: df=2, *X*^2^=0.037, p=0.98, Figure S4).

## 4 Discussion

*T. castaneum* is one of the rare species for which not only maternal but also paternal TGIP has been observed (Roth et al. 2010; Roth et al. 2018). It is therefore important to further study this phenomenon. One of the major open questions regarding paternal TGIP is, whether it is effective in more than one subsequent generation and can be considered to be multigenerational. Additionally, it is important to understand what the costs of TGIP are and if these are also transferred to later generations. We therefore carried out bacterial priming and challenge experiments across three generations using adult beetles.

We found that offspring of primed grandfathers survived a bacterial challenge significantly better than offspring of grandfathers, which had received a priming control injection. Thus, paternal TGIP is persistent for multiple generations at least until the F_2_ generation. Astonishingly, the survival advantage of the F_2_ generation was at a similar level as observed in previous experiments for the direct offspring (Eggert et al. 2014). We therefore did not see any dilution effect of this phenomenon over subsequent generation. Furthermore, we witnessed indirect costs, not of TGIP itself, but of the wounding procedure during the injection. These fitness costs became only visible after two generations, when the offspring of fathers from the injection control group sired significantly less offspring. In the present experiment, in contrast to previous studies (Roth et al. 2010; Eggert et al. 2014), were unable to detect a significant priming effect in the adult F_1_ offspring after paternal priming. This was likely due to an unusually high mortality in the injection control, maybe caused by a bacterial contamination in the injection buffer that was used for all treatments, thereby reducing a potential effect of priming.

Few studies have investigated the effects of TGIP beyond the first offspring generation. It has been shown that viral silencing agents derived from an RNAi response can be inherited non-genetically from either parent and passed on for several generations (Rechavi et al. 2011). In parthenogenetic *Artemia*, maternal exposure to bacteria provided the offspring with a survival benefit of bacterial infection for all three tested offspring generations (Norouzitallab et al. 2015). Multigenerational effects of paternal TGIP have been described in the pipefish, where due to male pregnancy contact between father and offspring is much more pronounced than in our system (Beemelmanns and Roth 2017). Although, we are as of today unaware of the mechanisms behind paternal TGIP against bacteria, we can assume that its multigenerational nature will strongly impact the evolution of resistance and tolerance, depending on the costs, benefits and specificity of TGIP and the prevalence of and therefore chances of repeated exposure to a parasite.

In the second part of this study, we investigated the transgenerational impact of immune priming via two different infection routes in larvae, for which within life stage immune priming has been previously demonstrated (Roth et al. 2009; Milutinović et al. 2014). Additionally to the survival after bacterial challenge, we monitored fitness costs of larval priming, becoming apparent as either directly reduced fertility or by slowing down developmental speed of the treated individual or its offspring. As any form of immunity, also immune priming comes at a cost for the organism (Sadd and Schmid-Hempel 2009; Schmid-Hempel 2005; Freitak et al. 2009). While in mosquitos a trade-off between immune priming and egg production has been observed (Contreras-Garduño et al. 2014), we did not find any effects of priming on fertility. Similar numbers of live offspring were produced across all treatments for both priming methods. But we estimated fertility only from a short 48h reproduction period and do not know how the immune priming might affect lifetime reproductive success. Also, we provided the beetle with *ad libitum* food throughout the experiment, whereas limiting resources can be necessary for uncovering trade-offs with immunity (Kutzer and Armitage 2016; Moret and Schmid-Hempel 2000).

However, the oral priming of larvae led to differential speed in their development. Larvae, which had received the priming control diet containing the supernatant from the *Bt407* culture reached pupation considerably faster and emerged as adults earlier. In contrast, the treatment with *Btt* did not lead to differential developmental time compared to the naïve larvae. The same effect was observed previously by Milutinović et al. (2014). It is possible that the supernatant from the *Bt407* control culture contained some nutrients that were transferred to the priming diet and helped the larvae to speed up their development. The supernatant from the treatment *Btt* culture might not contain these nutrients, due to differences in the bacteria. Alternatively, the necessity to mount an immune and priming response, brought on by the exposure to the priming diet might mitigate the potential effect of the additional nutrients.

In the offspring generation, development was strongly affected by parental larval treatment. Both, offspring from the *Btt* primed group and the *Bt407* priming control took longer to pupate and also emerge as adults. This is interesting because although *Bt407* does neither provide an immune priming (Milutinović et al. 2014) nor is able to kill larvae upon ingestion (Milutinović et al. 2013), larvae feeding on its spore supernatant still suffer these fitness costs. These results are in concordance with observations in the mealworm beetle, where maternal priming prolonged larval development, while paternal priming led to a reduction in larval body mass (Zanchi et al. 2011). For the injection priming, we only observed within generation effects on the development. Here the wounding by the injection was sufficient to cause the effect, because larval development was slowed down in the injection of heat-killed bacteria as well as in the injection control treatment compared to the naïve group. In the offspring generation, time until adult emergence was not affected by parental priming. So far, we have no data regarding the development until pupation in this case.

Increased developmental time during the larval and pupal stage can be considered a fitness cost. Longer time spend during the larval stage is costly as it increases several risks. During the larval stage the risk of infection is higher as only larvae can be infected orally with certain bacteria, including *Btt*. Also, there is a higher risk of cannibalism, which happens regularly among larva (Ichikawa and Kurauchi 2009) and at high densities smaller larvae might be less able to secure sufficient food (Koella and Boëte 2002). Therefore, prolonged development should decrease probability of survival and delay the start of reproduction. In this experiment we were unable to confirm within-generation immune priming for either of the two used infection methods. This can likely be attributed to the low overall mortality rates following the challenge, which is a problem occasionally encountered in such experiments (see also Tate et al. 2017). However, both within-generation priming methods have been shown to work consistently in our lab (Milutinović et al. 2014; Ferro et al. 2017; Futo et al. 2017).

We did not find any evidence of larval TGIP with the oral nor the injection protocol. For larval priming by septic wounding with a pricking needle, it was observed that TGIP in larvae only occurred in populations, which do not demonstrate within life stage immune priming (Khan et al. 2016), implying that they are incapable of developing and maintaining both forms of immune protection. As beetles from our population have repeatedly been shown to possess larval within life stage priming ability, this is a possible explanation for the absence of larval TGIP.

In conclusion, we observed that different ways of immune priming can have different effects on the next generation, depending on the life stage and route of priming. These effects might not always be beneficial, as parental treatment appeared to impact on offspring development, demonstrating potential costs of immune priming that are paid by the next generation. These would remain undetected if only the treated generation is studied. We therefore do not only need more studies on mechanisms behind the different routes of immune priming, but also more experimental research focusing on the evolutionary consequences of immune priming. This will help to clarify under which circumstances this ability is favored over the evolution of resistance or tolerance (Tidbury et al. 2012; Tate 2017). Such knowledge will be needed to advance methods of pest control, which strongly depend on the use of bacterial products, e.g. toxins from *B. thuringiensis*.

## 6 Author Contributions

All authors contributed to the conception and design of the study; NS, MPS, KF and NK performed the experimental work; NS performed the statistical analysis and drafted the manuscript. All authors contributed to manuscript revision and read and approved the submitted version.

## 7 Funding

We are grateful for the financial support provided to MPS and JK by the DFG grant KU 1929/8-1 within the DFG priority program 1819 “Rapid adaptations” and the DFG grant KU 1929/4-2 to KF and JK within the DFG priority program 1399 “Host-parasite coevolution.”

### 8 Acknowledgments

We wish to thank Kathrin Brüggemann und Anna Hübenthal for their help with the experiments. We thank Sina Flügge for providing us with drawings of *T. castaneum*.

## 10 Data Availability Statement

The datasets generated for this study can be found in the supplementary material.

